# Virulence and antimicrobial resistance features among clades of *Escherichia coli* ST131 strains causing community-acquired urinary tract infection in Rio de Janeiro, Brazil

**DOI:** 10.64898/2026.03.18.712724

**Authors:** Isadora Silva Barcellos, Thaís Costa da Conceição de Sousa, Eduardo Moreira de Castro, Juliana Jandre Sueiro de Souza Pereira, Adriana Lúcia Pires Ferreira, Káris Maria de Pinho Rodrigues, Beatriz Meurer Moreira, Karla Rodrigues Miranda

**Affiliations:** Universidade Federal do Rio de Janeiro, Instituto de Microbiologia Paulo de Goes, Rio de Janeiro, RJ, Brazil; Universidade Estácio de Sá, Faculdade de Medicina, Instituto de Educação Médica, Rio de Janeiro, RJ, Brazil; Diagnósticos da América S/A, DASA, Rio de Janeiro, RJ, Brazil

**Author notes:** Corresponding author: E-mail adress.

## Abstract

Urinary tract infection (UTI) is one of the most common community-acquired bacterial infections mainly caused by extraintestinal pathogenic *Escherichia coli* (ExPEC) strains. The high-risk *Escherichia coli* ST131 clone is a major global cause of this disease. The lineage rapid dissemination is associated to multidrug resistance (MDR), production of extended-spectrum beta-lactamase (ESBL), and multiple virulence-associated genes. Although we lack information about ExPEC high-risk clones in Latin America, we recently reported an increase in ST131 dissemination in Rio de Janeiro from 2015 to 2019. The present study aims to characterize virulence and resistance molecular and phenotypic features that may contribute to dissemination of *E. coli* ST131 in Rio de Janeiro, Brazil. We assessed a 133 *E. coli* ST131 strains collection obtained from urine of outpatients with suspected UTI, in 2019. We determined antimicrobial susceptibility, fluoroquinolones resistance genes, virulence factors associated genes and biofilm production of all strains and analyzed the frequencies by each clade or subclade. A higher incidence of women (92%) and elderly (65%) subjects was observed. Overall resistance to first- and second-line treatment for UTI antimicrobials ampicillin, ciprofloxacin and sulfamethoxazole-trimethoprim was detected in high rates (40%), with a major impact of subclade C2 strains that were resistant to almost all antimicrobials tested, 52% carry ESBL and 66% of strains harbor the *aac(6’)-Ib-cr* ciprpofloxacin resistance gene. Clade B and subclade C2 showed higher virulence scores among the other clades. They present unique virulence profiles characterized by the presence of *papGIII*, *sfa/focDE*, and especially *ibeA* genes in clade B, and the *afa/DrBC, papGII, hlyA*, *cnf1* genes in subclade C2. Over 50% of our strains are biofilm producers, characterized by weak (24%) and strong producers (32%). ESBL and MDR strains harbor mainly *papA, papGII, hlyA*, *cnf1* and *kpsMTII* genes that plays a key role in ST131 colonization. Subclade C1 is the major biofilm producer (78%), despite its lower virulence score. We also detected higher incidence of *papA* (27%), *hlyA* (19%) genes and the RPAI(*malX*) marker (84%) in biofilm producer strains with a statistical association of *sfa/focDE* gene (9%). We can infer that Clade C strains might be responsible for ST131 dissemination and persistence in Rio de Janeiro.

## 1. Introduction

Urinary tract infection (UTI) is one of the most frequent community-acquired bacterial infection and major source of outpatient health care visits, with over 400 million cases reported globally in 2019, including 240,000 fatalities (1). Extraintestinal pathogenic *Escherichia col*i (ExPEC) is responsible for over 80% of community-acquired UTIs and many healthcare-associated infections (2, 3). UTI requires antimicrobial therapy; therefore, it is not a surprise that ExPEC strains causing UTI are associated with the acquisition and spread of antimicrobial resistance (AMR) genes in community and hospital environments (3–5) presenting a severe global public health threat (6, 7). In 2019, the global burden of bacterial AMR study revealed that 1.27 million deaths were attributed to AMR bacteria that year worldwide. *E. coli* was the leading pathogen associated with AMR-related deaths, particularly in Latin America (8).

ExPEC high-risk clones are responsible for most infections and AMR spread (9, 10), with a major contribution of ST131, appearing in 90% of studies and increasingly dominating UTI cases since the 2000s (10). ST131 is subdivided into clades and subclades A, B, C1, C1-M27 and C2. Clade C is the most widely disseminated, representing 80% of global ST131 isolates (11–13). This dominance has been attributed to multidrug resistance (MDR), especially fluoroquinolone resistance and CTX-M production in subclade C2.

ExPEC pathogenicity is likely related to the expression of a set of virulence genes. In recent years, studies have indicated that ST131 Afa/Dr adhesin and CNF1 toxin aid host gut colonization, while the PapGII adhesin interacts with resistance genes, potentially fueling clade evolution and dominance (14–16). In addition, biofilm formation is a crucial virulence factor (VF) of uropathogenic *E. coli* (UPEC) that can be one of the key factors for the host’s colonization, and recurrence of UTIs. UPEC biofilm can lead to the formation of “*intracellular bacterial communitites*” (IBC) and reservoirs (2) and protects bacteria against harmful environments, antimicrobial agents, and the host’s immune system (17).

Despite the high ST131prevalence, comprehensive data about the epidemiological profile and biomarkers of high-risk *E. coli* clones are missing for South America (10, 18). In 2019, data from the Global Burden Disease (GBD) study showed that Tropical Latin America had the highest age-standardized incidence rates of UTI per 100,000 population (1, 19). Although few studies have reported high-risk ExPEC clones isolated from UTI in Brazil, ST131 was predominant in all studied populations (20–22). We previously showed a significant increase in the occurrence of ST131 in Rio de Janeiro, from 3% in 2005 to 9% in 2015, and 14% in 2019 (23). However, the virulence potential of such strains from Brazil remains poorly characterized. This study aims to analyze the molecular and phenotypic virulence and resistance profiles of *E. coli* ST131 clades to identify potential factors driving their evolution and clinical impact.

## 2. Methods

### 2.1. Subjects, clinical specimens, and bacterial isolates

We studied a collection of 133 *E. coli* strains isolated from urine specimens of outpatients with suspected UTI in Rio de Janeiro state in July 2019. The following procedures described were performed to all isolates. Strains were previously classified as ST131, typed for clades/subclades and ESBL production was established. Regarding clades distribution, our collection comprises Clade A: 21 strains, Clade B: 35 strains, subclade C1: 36 strains, subclade C1-M27: 7 strains and subclade C2: 33 strains. We couldn’t classify one strain into clades, named non-typable (NT) (23).

### 2.2. Antimicrobial susceptibility testing

Antimicrobial susceptibility was determined by disk diffusion method according to Clinical and Laboratory Standards Institute (CLSI) guidelines. Susceptibility profiles were interpreted as stated by CLSI 2025. The test was performed for the following drugs: amikacin, amoxicillin-clavulanate, ampicillin, aztreonam, cefepime, cefotaxime, cefoxitin, ceftazidime, ceftriaxone, ciprofloxacin, ertapenem, gentamicin, nitrofurantoin, and sulfamethoxazole-trimethoprim. The ATCC® 25922 strain was used as control. Multidrug resistance (MDR) was defined as non-susceptibility to one or more agents in three or more antimicrobial classes, as suggested (24).

### 2.3. Molecular detection of resistance and virulence encoding genes

Multiplex PCR was performed to detect the plasmid-mediated fluoroquinolone resistance (PMQR) genes *qnr*, *qepA* and *aac(6’)-Ib* (25–28). The final product of *aac(6’)-Ib* positive strains was purified with ExoProStar 1-Step (Cytiva, EUA) and then sequenced by SeqStudioTM (AppliedBiosystems – ThermoFisher). We determined the *aac(6’)-Ib-cr* variant sequences with BioEdit v7.2.5 analyses and comparisons by BLAST with data available at the NCBI database (NG_047292.1, NG_052086.1 and NG_067968.1 were used as reference), considering 99-100% of identity and 90% of coverage. Virulence-associated genes commonly found in UPEC strains *papAH, papGI, papGII, papGIII, sfa/focDE, afa/DR, cnf1, hlyA, kpsMTII, fyuA, iutA, malX, traT* and *ibeA* were detected by multiplex PCR, according to previous published protocols (29–36).

### 2.4. Biofilm production

Briefly, strains were cultured in TSA medium and grown overnight in aerobic atmosphere at 35 ± 2 °C. One colony was transferred to 3 mL of tryptic soy broth (TSB) medium and then incubated overnight at 35± 2 °C in a stationary condition. Following, 200 μl of 1:100 bacterial dilution was inoculated into sterile microtiter wells, in quadruplicate. The sterile medium was used as negative control. *E. coli* control strains were ST131 EC15_02 (negative), and EAEC 042 (positive). After incubation of inoculated plates for 24 h at 35± 2 °C, planktonic cells were aspirated, and the biofilm was fixed by dry heat at 60 °C for 1 hour. Wells were washed twice with phosphate buffered saline (PBS; pH7) and dried for 10 min at room temperature. The biofilm was stained with 200 μl of crystal violet (0.1%) for 15 min, and wells were washed twice with PBS. Finally, 200 μl of an 80% ethanol and 20% acetone solution were added in the wells and plates were incubated for 30 min at room temperature. Optical density (OD) measurements were generated by an ELISA reader at 590 nm. Strains were classified as strong, weak and non-biofilm producers, according to the criteria previous published (37). The experiment was performed in triplicate.

### 2.5. Statistical analysis

Virulence and PMQR encoding genes and AMR frequencies are described as counts and percentages. Clades and subclades frequencies, AMR, MDR and ESBL frequencies were obtained from a previous study (23). Univariate analyses of the distribution of gender, resistance to each agent, MDR and ESBL production, PMQR and virulence encoding genes by clades and biofilm production were performed with the Chi-Square or Fisher’s exact tests. Virulence scores were calculated. For each isolate we assessed the sum of *pap* (presence of *papAH* and/or *papG* allele II and/or *papG* allele III was counted as 1 unit) and all the other VFs detected. Means were calculated and analyzed with Mann–Whitney U test (38). We used SPSS® Statistics V.23 and GraphPad Prism V. 8.02 for statistical analysis and graphic generation. We defined significance for two-tailed p-value *<* 0.05.

### 2.6. Ethics

The study was approved by the Human Research Ethics Committee of Universidade Federal do Rio de Janeiro (CAEE 59953322.9.0000.5257).

## 3. Results

From the collection of 133 *E. coli* ST131 strains, over 90% of the individuals were women, 65% (N=87) were over 59 years old, and among them, a slightly higher incidence of subclade C2 isolates was observed (**Table 1**).

**Table 1:**
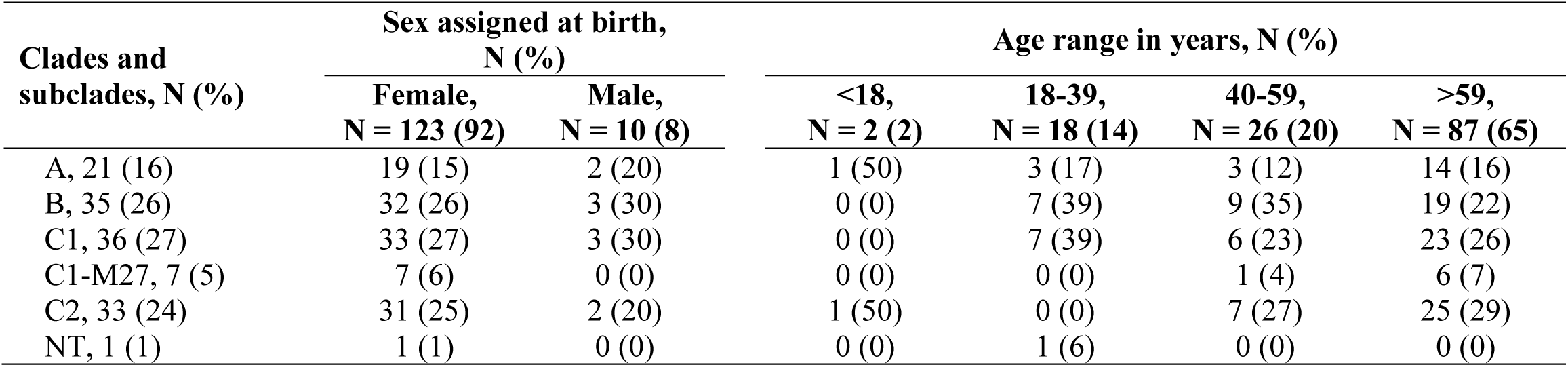
Distribution 133 *E. coli* ST131 isolates in clades by sex and age range. NT: non-typable

| Clades and<br>subclades, N (%) | Sex assigned at birth,<br>N (%) |  | Age range in years, N (%) |  |  |  |
| --- | --- | --- | --- | --- | --- | --- |
|  | Female,<br>N = 123 (92) | Male,<br>N = 10 (8) | <18,<br>N = 2 (2) | 18-39,<br>N = 18 (14) | 40-59,<br>N = 26 (20) | >59,<br>N = 87 (65) |
| A, 21 (16) | 19 (15) | 2 (20) | 1 (50) | 3 (17) | 3 (12) | 14 (16) |
| B, 35 (26) | 32 (26) | 3 (30) | 0 (0) | 7 (39) | 9 (35) | 19 (22) |
| C1, 36 (27) | 33 (27) | 3 (30) | 0 (0) | 7 (39) | 6 (23) | 23 (26) |
| C1-M27, 7 (5) | 7 (6) | 0 (0) | 0 (0) | 0 (0) | 1 (4) | 6 (7) |
| C2, 33 (24) | 31 (25) | 2 (20) | 1 (50) | 0 (0) | 7 (27) | 25 (29) |
| NT, 1 (1) | 1 (1) | 0 (0) | 0 (0) | 1 (6) | 0 (0) | 0 (0) |

Higher incidence of resistance was noticed to first- and second-line treatment for UTI antimicrobials: ampicillin, ciprofloxacin (both >70%), and sulfamethoxazole-trimethoprim (>40%) (**Table 2)**. Unlikely, Clade A impacted on overall ampicillin resistance rate, as it was the highest among clades (95%; p=0.03). Clade C was the major contributor to ciprofloxacin resistance: C1: 86%, p=0.04 and C2: 100%, p<0.001. Of the 16 antimicrobials tested, subclade C2 showed higher resistance to 11 antimicrobials compared to the other clades (p<0.05), including first line antimicrobials as nitrofurantoin and thrimethoprim/sulfamethoxazole. This led to 79% (p=0.0002) of MDR and 52% (p<0.001) of ESBL production in subclade C2 isolates. Since all subclade C2 strains are ciprofloxacin resistant, the presence of the PMQR genes *aac(6’)-Ib-cr* (66%, p<0.0001) and *qnrB* (27%, p=0.02) were higher compared to the other clades **(Table 2)**.

**Table 2:**
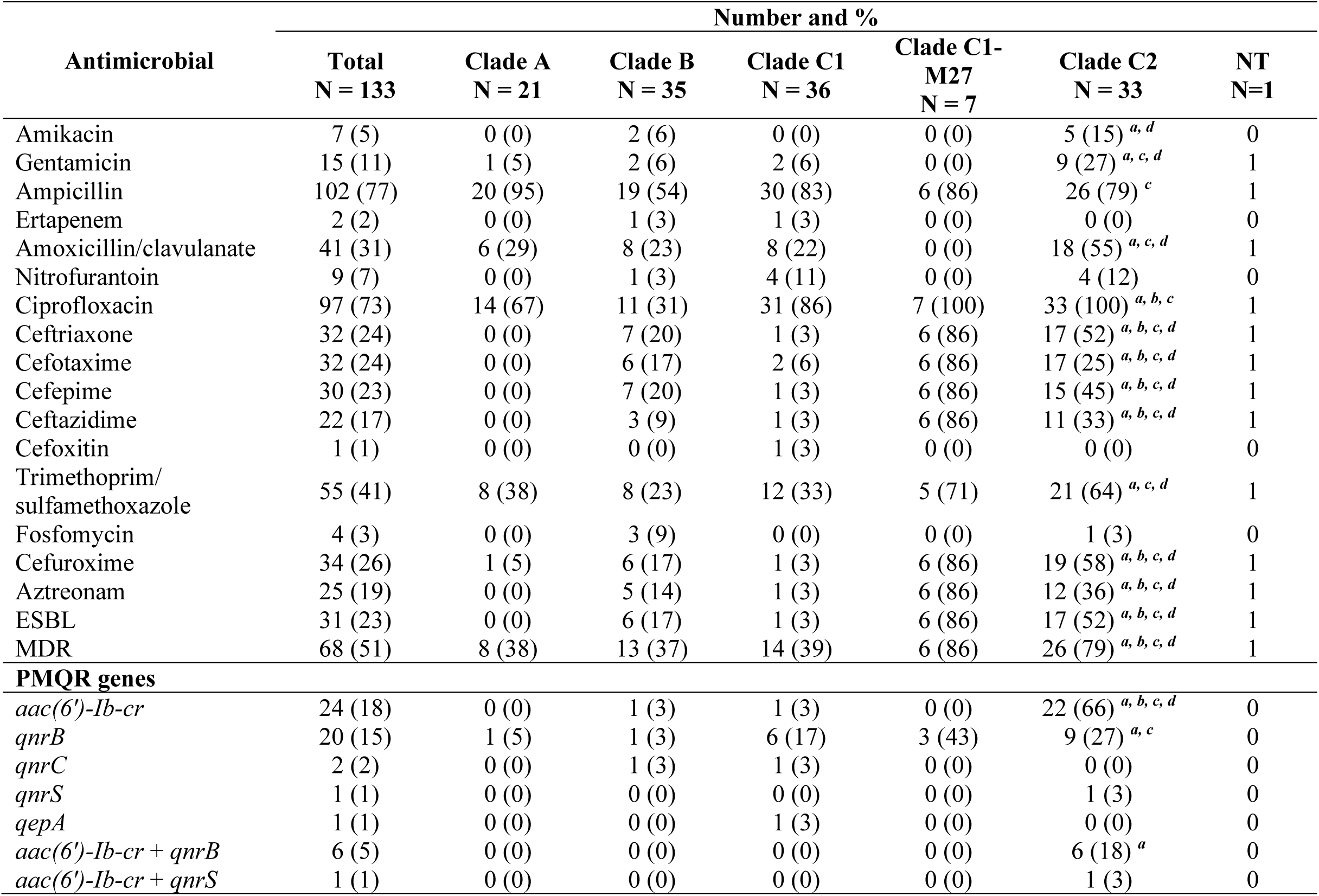
Distribution of antimicrobial non-susceptible *E. coli* ST131 isolates and PMQR genes by clade and subclade. NT: non-typable; ESBL: Extended spectrum betalactamase; MDR: multidrug resistance; PMQR: plasmid mediated quinolone resistance; superscript letters indicate a significant difference (p<0.05), compared with *^a^*other clades; *^b^*clade A; *^c^*clade B; *^d^*clade C1.

Subclade C2 showed higher resistance to ciprofloxacin, ceftriaxone, cefotaxime, ceftazidime, cefuroxime, and aztreonam, compared to Clade A (p<0.001) and presented non-susceptibility to most antimicrobials compared to Clade B and subclade C1 (p<0.05), except for ampicillin in subclade C1 (**Table 2**). We also evaluated the frequency of PMQR genes in ESBL-producing samples and MDR strains to infer the resistance burden. The *aac(6’)-Ib-cr* gene (55%, p<0.001) and *qnrB* gene (26%, p=0.055) were detected more frequently in ESBL-producing strains. In MDR strains, only the *aac(6’)-Ib-cr* gene (32%, p<0.001) was associated.

No statistically significant differences were observed in resistance rates, ESBL production, or MDR classification between sex or age groups.

We investigated the main virulence genes detected in ExPEC strains. The predominant genes in this collection were *fyuA* which encodes a siderophore receptor, *iutA*, which encodes the ferric aerobactin receptor IutA, and the pathogenicity island marker RPai(*malX*) (**Table 3**). We also calculated the virulence score, *i.e*., the average sum of virulence genes detected in each clade. The highest virulence scores were observed in Clade B and subclade C2 (208; 5.9 and 206; 6.2) though subclade C1 showed the lowest (135; 3.75) **(Figure 1)**.

**Figure 1:**
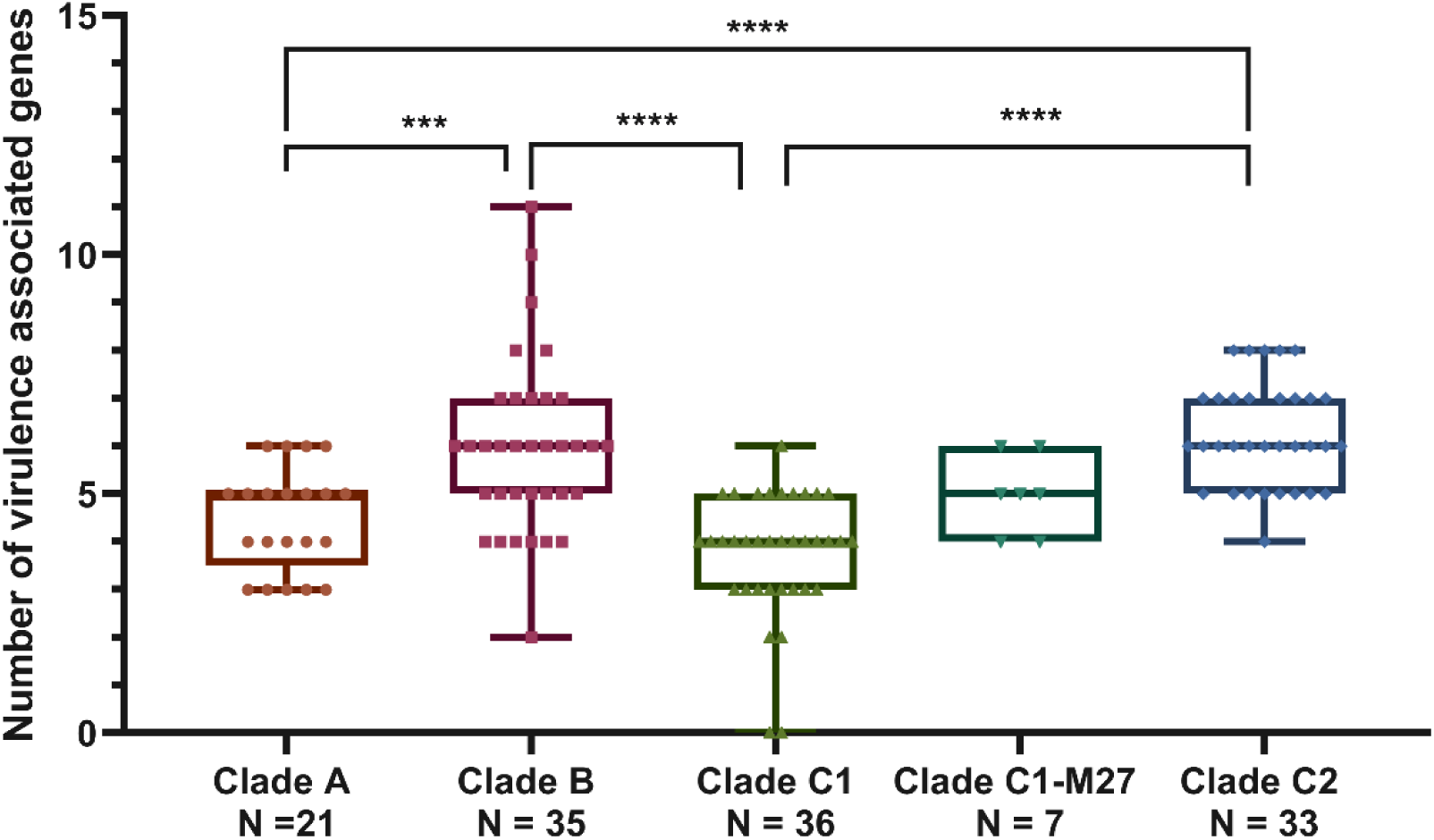
Virulence score of each clade. Represents the sum and mean of virulence encoding genes detected in each clade. ***: p = 0.0007, ****: p <0.0001

**Table 3:**
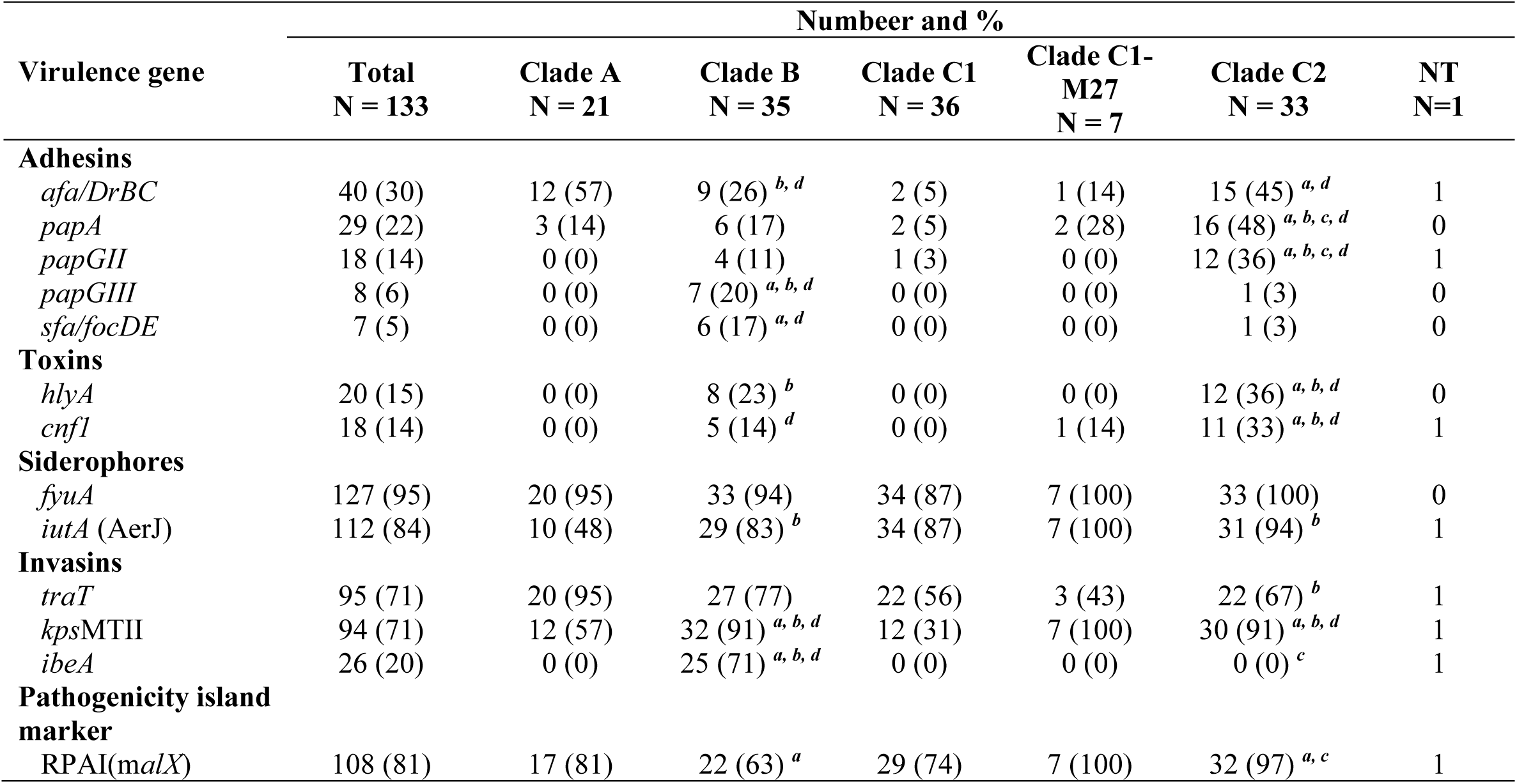
Distribution of virulence factors encoding genes by clades. NT: non-typable; superscript letters indicate a significant difference indicate a significant difference (p<0.05), compared with *^a^*other clades; *^b^*clade A; *^c^*clade B; *^d^*clade C1.

|  | Number and % |  |  |  |  |  |  |
| --- | --- | --- | --- | --- | --- | --- | --- |
| Virulence gene | Total<br>N = 133 | Clade A<br>N = 21 | Clade B<br>N = 35 | Clade C1<br>N = 36 | Clade C1-M27<br>N = 7 | Clade C2<br>N = 33 | NT<br>N=1 |
| **Adhesins** |  |  |  |  |  |  |  |
| *afa/DrBC* | 40 (30) | 12 (57) | 9 (26) <sup>b, d</sup> | 2 (5) | 1 (14) | 15 (45) <sup>a, d</sup> | 1 |
| *papA* | 29 (22) | 3 (14) | 6 (17) | 2 (5) | 2 (28) | 16 (48) <sup>a, b, c, d</sup> | 0 |
| *papGII* | 18 (14) | 0 (0) | 4 (11) | 1 (3) | 0 (0) | 12 (36) <sup>a, b, c, d</sup> | 1 |
| *papGIII* | 8 (6) | 0 (0) | 7 (20) <sup>a, b, d</sup> | 0 (0) | 0 (0) | 1 (3) | 0 |
| *sfa/focDE* | 7 (5) | 0 (0) | 6 (17) <sup>a, d</sup> | 0 (0) | 0 (0) | 1 (3) | 0 |
| **Toxins** |  |  |  |  |  |  |  |
| *hlyA* | 20 (15) | 0 (0) | 8 (23) <sup>b</sup> | 0 (0) | 0 (0) | 12 (36) <sup>a, b, d</sup> | 0 |
| *cnfI* | 18 (14) | 0 (0) | 5 (14) <sup>d</sup> | 0 (0) | 1 (14) | 11 (33) <sup>a, b, d</sup> | 1 |
| **Siderophores** |  |  |  |  |  |  |  |
| *fyuA* | 127 (95) | 20 (95) | 33 (94) | 34 (87) | 7 (100) | 33 (100) | 0 |
| *iutA* (AerJ) | 112 (84) | 10 (48) | 29 (83) <sup>b</sup> | 34 (87) | 7 (100) | 31 (94) <sup>b</sup> | 1 |
| **Invasins** |  |  |  |  |  |  |  |
| *traT* | 95 (71) | 20 (95) | 27 (77) | 22 (56) | 3 (43) | 22 (67) <sup>b</sup> | 1 |
| *kpsMTII* | 94 (71) | 12 (57) | 32 (91) <sup>a, b, d</sup> | 12 (31) | 7 (100) | 30 (91) <sup>a, b, d</sup> | 1 |
| *ibeA* | 26 (20) | 0 (0) | 25 (71) <sup>a, b, d</sup> | 0 (0) | 0 (0) | 0 (0) <sup>c</sup> | 1 |
| **Pathogenicity island marker** |  |  |  |  |  |  |  |
| RPAI(*malX*) | 108 (81) | 17 (81) | 22 (63) <sup>a</sup> | 29 (74) | 7 (100) | 32 (97) <sup>a, c</sup> | 1 |

Comparing clade B with the remaining clades, *ibeA* gene, which encodes an invasin of brain endothelial cells, was the only VF found in a single clade, suggesting a virulence signature of clade B, in this collection. Other VFs as *papGIII* (p<0.001), *sfa/focDE* (p=0.001), *kpsMTII* (p=0.002), and *ibeA* (p<0.0001) were predominant, while the RPAI(*malX*) marker was less detected (63%, p=0.001). The gene encoding the Afa/Dr adhesin was the only one with lower detection rate in Clade B, compared to A (p=0.02). Subclade C2 also shows a unique virulence profile as the genes *afa/DrBC* (p=0.03), *papAH*, *papGII*, *hlyA*, *cnf*, *kpsMTII* (p<0.001), and RPAI(*malX*) (p<0.01) were the predominant VFs compared to the remaining strains. Comparing subclade C2 with Clade A, six genes were more frequent in this clade (p<0.05), while the gene *traT* was associated with Clade A (p=0.01) (**Table 3**).

We assessed whether there is a relationship between the detection of virulence genes and the sex or age range of individuals and observed that the *afa/DrBC* gene was more frequent in women than in men (33%; p=0.03). The pathogenicity island marker RPAI(*malX*) was predominant in individuals <59 years compared to other ages (87%, p=0.01). However, it is noteworthy that women and individuals over 59 years of age are the predominant investigated population.

To evaluate co-existence of AMR and VFs we analyzed the virulence profile according to ESBL production and the MDR phenotype (**Figure 2**). The genes encoding the adhesins *papAH*, *papGII*, the toxins *hlyA* and *cnf1*, and the invasin *kpsMTII* are predominant in ESBL-producing strains compared to non-producing strains (**Figure 2A**). Similarly, the virulence score of ESBL-producing strains was significantly higher (189; mean 6.1; p<0.0001) (**Figure 2C**). In MDR strains, the genes *papAH, papGII*, and *hlyA* were more frequent, and the gene *ibeA* was associated with non-MDR strains (**Figure 2B**).

**Figure 2:**
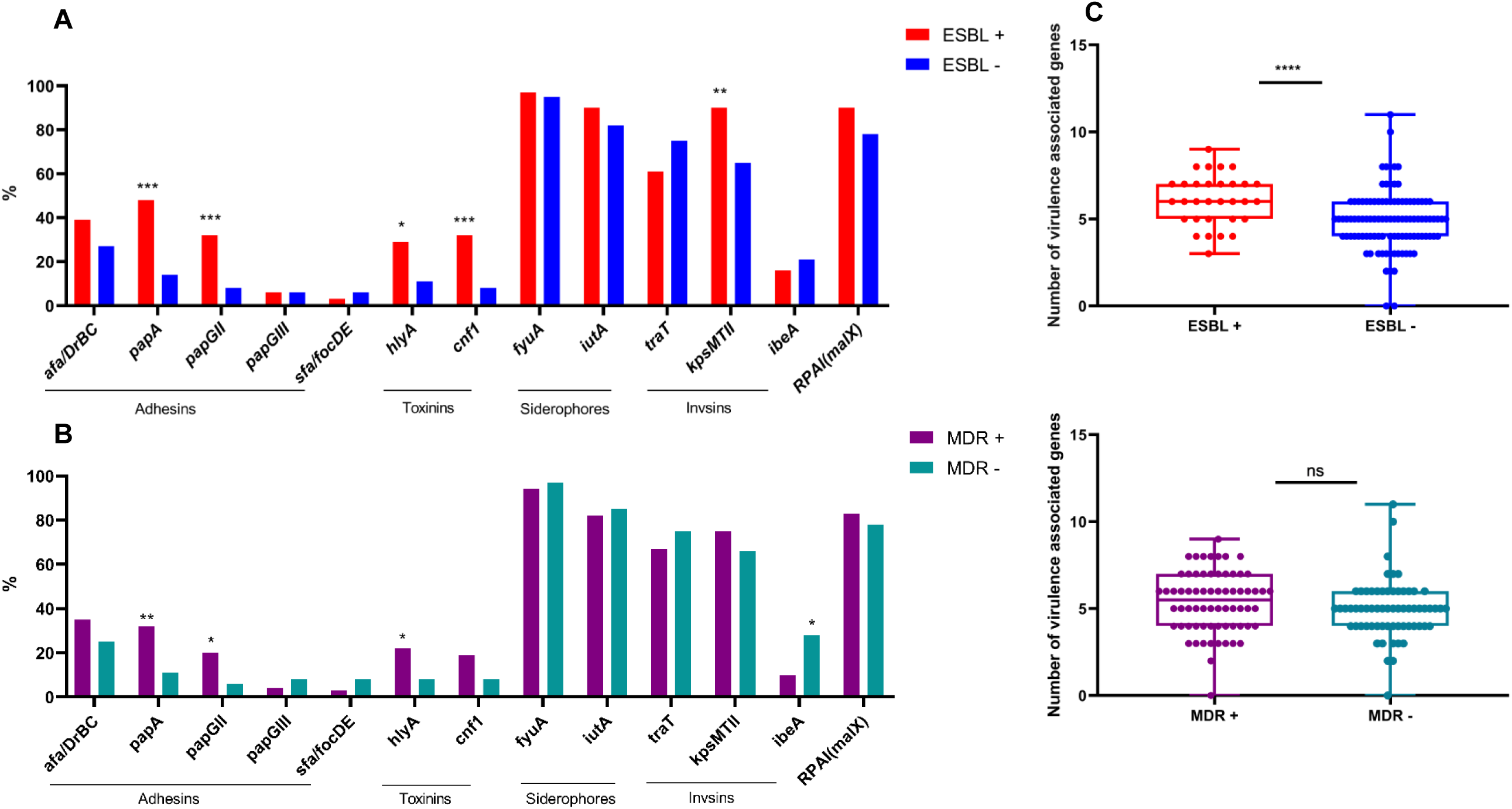
Virulence profile by resistance phenotype. Frequency of virulence factors encoding genes in A) ESBL producers and non-ESBL producers strains and B) MDR strains and non-MDR strains; C) Virulence score of ESBL and MDR strains. * p<0.05; ** p<0.01; *** p<0.001; **** p<0.0001.

Over 50% of strains could form biofilm in which 24% (N=32) are weak producers and 34% (N=45) are strong producers (**Table 4**). Clade C seems to be an important contributor to biofilm formation in ST131 as C1 was the highest producer (N=28, 78%), followed by subclades C1-M27 (N=4, 57%) and C2 (55%, N=18) and the production by subclade C1 was statistically significantly higher compared to all clades (p=0.005) (**Table 4 and Figure 3**). Concerning the intensity of biofilm production, almost all clades showed a higher incidence of strong production, except for subclade C1 that was equally weak and strong producer **(Table 4).** We observed a progressive increase incidence of VFs papA, hlyA, sfa/focDE and RPAI(malX) among non-biofilm producer, weak and strong biofilm producer strains. Yet only *sfa/focDE* gene was associated with biofilm-producing samples (p=0.02) **(Table 5).** No statistically significant differences were observed in virulence genes between strong and weak biofilm producers, nor in the resistance phenotype with the biofilm production classification.

**Figure 3:**
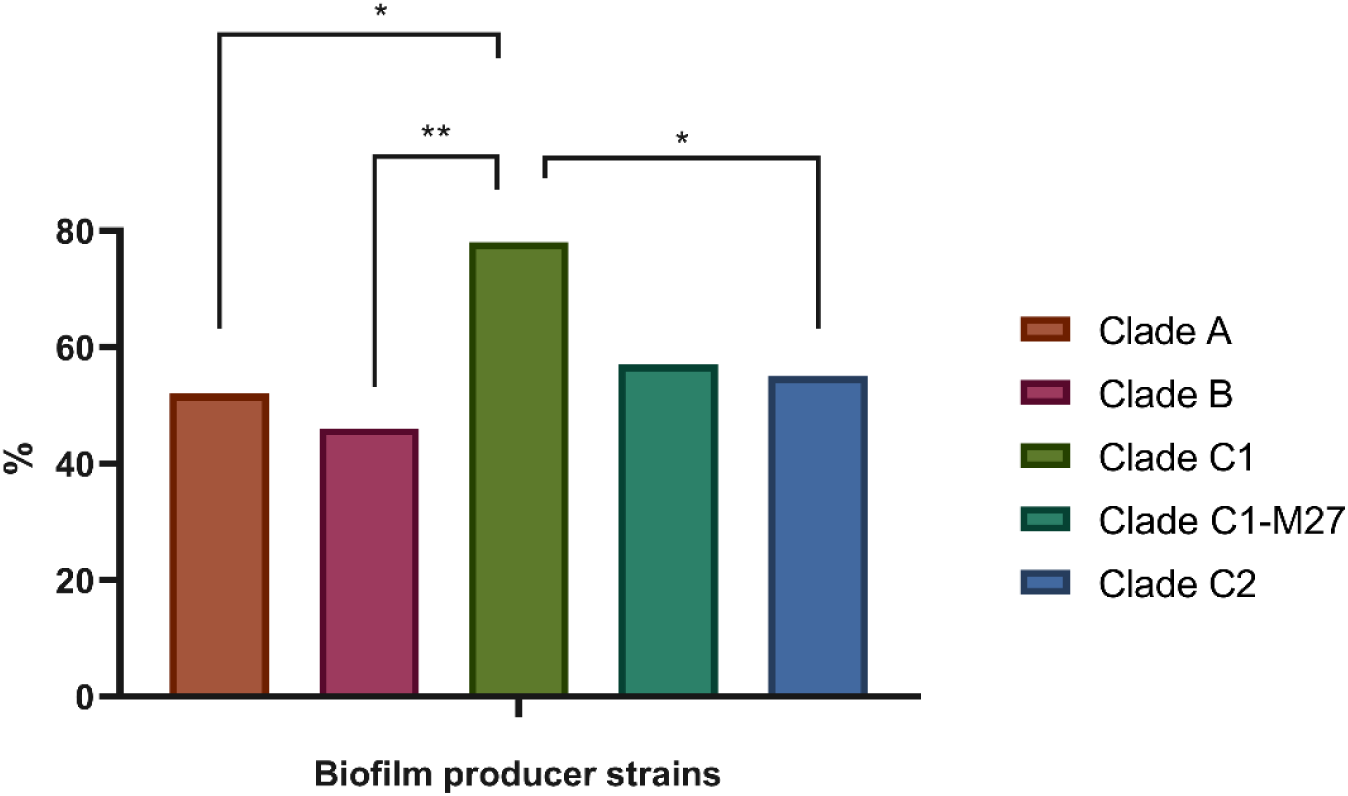
Frequency of biofilm production by clades. Statistically significant results comparing C1 to Clade A or C2 *: p <0.05 and C1 to Clade B **: p = 0.0054.

**Table 4:** Biofilm production classification among clades. *: p values indicate a significant difference (p=0.004664) compared to other clades.

| Biofilm production classification | Number and % |  |  |  |  |  | NT<br>N=1 (%) |
| --- | --- | --- | --- | --- | --- | --- | --- |
|  | Total<br>N = 133 (%) | Clade A<br>N = 21 (%) | Clade B<br>N = 35 (%) | Clade C1<br>N = 36 (%) | Clade C1-M27<br>N = 7 (%) | Clade C2<br>N = 33 (%) |  |
| Non-producer | 56 (42) | 10 (48) | 19 (54) | 8 (22) | 3 (43) | 15 (45) | 1 (100) |
| Producers | 77 (58) | 11 (52) | 16 (46) | 28 (78) * | 4 (57) | 18 (55) | 0 (0) |
| Weak biofilm | 32 (24) | 5 (24) | 5 (14) | 14 (39) | 1 (14) | 7 (21) | 0 (0) |
| Strong biofilm | 45 (34) | 6 (29) | 11 (31) | 14 (39) | 3 (43) | 11 (33) | 0 (0) |

**Table 5:** Distribution of virulence factors encoding genes detection by biofilm classification. *: p = 0,02 for comparison between non-biofilm producers *vs* biofilm producers.

| Virulence genes | Number and % |  |  |  |
| --- | --- | --- | --- | --- |
|  | Non-biofilm<br>producer<br>N = 56 (%) | All biofilm<br>producers<br>N = 77 (%) | Biofilm producer |  |
|  |  |  | Weak biofilm<br>N = 32 (%) | Strong<br>biofilm<br>N = 45 (%) |
| **Adhesins** |  |  |  |  |
| *afa/DrBC* | 21 (38) | 19 (25) | 9 (28) | 10 (22) |
| *papA* | 8 (14) | 21 (27) | 6 (19) | 15 (33) |
| *papGII* | 7 (13) | 11 (14) | 5 (16) | 6 (13) |
| *papGIII* | 1 (2) | 7 (9) | 3 (9) | 4 (9) |
| *sfa/focDE* | 0 (0) | 7 (9) * | 2 (9) | 5 (11) |
| **Toxins** |  |  |  |  |
| *hlyA* | 5 (9) | 15 (19) | 5 (16) | 10 (22) |
| *cnfI* | 4 (7) | 14 (18) | 7 (22) | 7 (16) |
| **Siderophores** |  |  |  |  |
| *fyuA* | 53 (95) | 74 (96) | 30 (96) | 44 (98) |
| *iutA* (AerJ) | 45 (80) | 67 (87) | 28 (87) | 39 (87) |
| **Invasins** |  |  |  |  |
| *traT* | 41 (73) | 54 (70) | 22 (59) | 32 (71) |
| *kpsMTII* | 43 (77) | 51 (66) | 19 (59) | 32 (71) |
| *ibeA* | 12 (21) | 14 (18) | 5 (16) | 9 (20) |
| **Pathogenicity**<br>**island marker** |  |  |  |  |
| RPAI(*malX*) | 43 (77) | 65 (84) | 25 (78) | 40 (89) |

## 4. Discussion

We report a high incidence of strains isolated from women and individuals older than 59 years. Although UTIs can affect anyone, they are more frequent in women (2) and are considered the most sex-biased disease due to lifetime disparities. Approximately 50% of adult women experience at least one UTI episode (1), whereas men have a 20–40 times lower risk (39, 40). However, prevalence increases in elderly men, mainly due to catheter use, and decreases in women (40). In our collection, 70% of male isolates were from individuals over 59 years. The higher prevalence of UTI in women is largely attributed to urinary tract anatomy, although other biological variations, as hormonal differences, may also play a role in this outcome (40). According to research based on GBD 2019, global increases in UTI incidence are linked to population aging and longer life expectancy (1, 19). Thus, it remains unclear whether susceptibility to *E. coli* ST131 is driven by pathogenicity or host/environmental features, highlighting the need for more and new UTI model studies.

The success of *E. coli* ST131 is still not fully understood, but multidrug-resistant (MDR) strains are likely key contributors (41). This lineage is characterized by CTX-M ESBL production and fluoroquinolone resistance. *E. coli* ST131 accounts for 70% of ESBL-producing and 78% of fluoroquinolone-resistant isolates in studies from India, the USA, and New Zealand (42–44). In our previous study on 2019 *E. coli* UTI collection, ST131 showed higher antimicrobial resistance (AMR) than other high-risk clones, especially to ampicillin, ciprofloxacin, and showed higher ESBL production. Although male isolates contributed to overall AMR (23), in the present study no significant sex-based differences were observed within ST131 AMR, suggesting this lineage did not drive that pattern.

In the present study, we further showed that ST131 sublineages differ markedly in frequency and AMR impact. Not surprisingly, subclade C2 was the main contributor to overall AMR, showing the highest resistance to 11 of 16 antimicrobials and increased ESBL production, among other clades. High ciprofloxacin resistance was linked to Clade C, with 100% resistance in C1-M27 and C2, and 86% in C1. Despite not being first-line therapy recommended by Infectious Diseases Society of America (IDSA) (45), fluoroquinolones remain widely prescribed for patients in several countries, including in Brazil (46–47). In fact, the Brazil- Global Antimicrobial Resistance Surveillance System (BR-GLASS) reported 32% ciprofloxacin resistance in community-acquired *E. coli* in Brazil, which was above the global average of 28% (48). We also emphasize the concerning increase resistance of first line antimicrobials in subclade C2, which lead to ST131 overall resistance to nitrofurantoin and trimethoprim/sulfamethoxazole in this collection. Clades A and B are generally more susceptible to antimicrobials, though their global prevalence may be underestimated due to selection bias (49). In our study, Clade B showed resistance to key antimicrobials, including amoxicillin/clavulanate, ampicillin, ciprofloxacin, and trimethoprim/sulfamethoxazole, although these rates aren’t statistically significant. Few studies assess AMR at subclade level, though findings from Iran support our results (50, 51). Finally, we observed a 66% incidence of the *aac(6’)-Ib-cr* gene in subclade C2, a variant of the aminoglycoside acetyltransferase that conferres resistance to ciprofloxacin and norfloxacin (52). This aligns with a recent report of 72% prevalence in subclade C2 and co-occurrence with *bla*_CTX-M-15_ (83%) (53), supporting our finding of 55% in ESBL strains. This gene may enhance survival under ciprofloxacin exposure, contributing to population-wide resistance. In addition to chromosomal resistance to quinolones, a dual resistance mechanism may contribute to ST131 persistence in host and its global dissemination. However, not all ciprofloxacin resistant strains in our collection harbor it, indicating the need to investigate chromosomal mutations (*gyrA* and *parC*).

Another explanation for ST131 dissemination is its pathogenicity based on VF content. It is considered the most virulent lineage within phylogroup B2 (54), with higher virulence scores than other high-risk clones (55). The most frequent VFs in our isolates were related to iron acquisition and pathogenicity islands, consistent with studies from Brazil (22, 56), Iran (50, 51), and Spain (38, 57, 58). Subclades differed in virulence profiles, with clade B and subclade C2 showing the highest scores consistent to other studies (50, 57), presenting individual signatures, such as the *papGIII*, *sfa/focDE*, and especially *ibeA* genes in clade B, and the *afa/DrBC, papGII, hlyA*, *cnf1* genes in subclade C2.

Subclade C2 has been consistently reported as more virulent than other C subclades in Spain (57–59). Clades A and subclades C1 and C1-M27 can be considered less virulent. Clade A was associated with Afa/DrBC adhesin and TraT invasin, but not KpsMTII, IutA, or Sat VFs, as proposed by other authors (50, 56,57). Clade B hallmark is the presence of the invasion of brain endothelial cells encoding gene *ibeA*, reported before (16, 57–60). In fact, if we remove *ibeA* gene from the clade B, its virulence score will reduce significantly from 208; 5.9 to 183; 5.2 and also will become statistically significant compared to subclade C2, reinforcing the importance of this gene to this clade. As far as we know, there is no study that investigate its role in Clade B dissemination, colonization, persistence in the host or infection process. Recently it was found that *ibeA* is present in a ST131 cluster related to avian and human hosts (60). Recent studies suggest C2 drives global ST131 dissemination. The *afaC* gene in 41% of global genome C2 strains enhances renal adhesion (14), while *papGII* is associated with pyelonephritis and AMR (15). CNF1 toxin, present in 76% of global C2 genomes, contributes to gut colonization but not UTI (16). Despite these findings and hypotheses, more studies are needed to compare the virulent potential of E. coli ST131 subclades with different molecular virulence markers in gut colonization or UTI. Given that multiple ST131 strains can colonize an individual (Forde et al., 2019), it is vital to identify which strain confers a competitive advantage.

We found that adhesins (*papAH, papGII*), toxins (*hlyA, cnf1*), and *kpsMTII* VFs were more frequent in ESBL strains, resulting in higher virulence scores. Notably, this result may be biased since 50% of subclade C2 are ESBL producers. Therefore, this finding likely reflects VF frequency in subclade C2. Similarly, *papAH*, *papGII*, and *hlyA* were associated with MDR strains, while *ibeA* was linked to non-MDR strains, likely due to clade distribution. The possible bias is due to the 79% MDR C2 strains and the Clade B strains non-MDR, that harbors the ibeA gene. Despite potential bias, the coexistence of AMR and VFs in community strains highlights their role in ST131 spread and treatment difficulty. The emergence of papGII+ C2 lineages, with integrated blaCTX-M-15 and plasmid replacement, may drive ST131 evolution (15), consistent with findings from Tunisia (61).

UTIs are facilitated by biofilm formation, which enhances bacterial survival and recurrence (17). Although widely studied in UPEC, biofilm formation in ST131 remains underexplored. In our study, 58% of strains formed biofilms, consistent with reports from outpatients with UTI (51), bloodstream infection (62), and UTI from children with malignant tumors (63). Interestingly, subclade C1, despite the lowest virulence score, was the main biofilm producer. Previous studies associated biofilm formation in ExPEC to several VFs such as *fimH, afa, kpsMTII, papA, cnf1, sfa/focDE, hlyA* (62, 64, 65). In this study we observed a progressive incidence of *papA*, *hlyA* genes and the RPAI(*malX*) marker among non-biofilm producers, weak and strong biofilm producer strains, although only *sfa/focDE* gene was statistically associated to biofilm production. This gene encodes fimbriae involved in adhesion to urinary epithelium; however, it was absent in C1 and present mainly in clade B. Further investigations of relative expression with transcriptomics or RT-qPCR are necessary to investigate exactly which gene is more expressed in biofilms.

In conclusion, ST131 strains causing community-acquired UTIs in Rio de Janeiro share global AMR and virulence features. Taking together, we can infer that in this collection, Clade C, especially subclade C2, appears central to local dissemination as it plays an important role as reservoir of resistance to the main antimicrobials used in clinical practice, potentially disseminating resistance genes that often coexist in one single strain due to high AMR. Furthermore, we observed that, MDR and ESBL-producing strains, accumulate important VF that have already been reported in the literature as the key contributors to the colonization of individuals with ST131 strains. Meanwhile subclade C1 contributes through biofilm formation that could contribute to ST131 persistence and increase in Rio de Janeiro. These findings indicate that virulence depends not only on VF presence but also on expression, adaptation, and metabolism. The diversity among clades and subclades helps explain ST131’s rapid global spread. Future surveillance should focus on clade/subclade-specific characteristics to avoid bias and improve control strategies for *E. coli* ST131 dissemination. Genomic and *in vivo* UTI model studies are essential to refine knowledge in clades epidemiology and virulent potential. Understanding the factors driving the expansion of resistant and virulent clones like *E. coli* ST131, including biofilm formation, is crucial for controlling dissemination, reducing infection severity, and managing recurrent UTIs, especially in low- and middle-income countries where data on ExPEC high-risk clones remain limited.

## Acknowledgements

This study was supported by Instituto Nacional de Pesquisa em Resistência aos Antimicrobiana (INPRA) of Conselho Nacional de Desenvolvimento Científico e Tecnológico (CNPq) and Rede de Resistência aos Antimicrobianos (REDES) under Fundação Carlos Chagas Filho de Amparo à Pesquisa do Estado do Rio de Janeiro (FAPERJ).

